# Assessing genomic reproducibility of read alignment tools

**DOI:** 10.1101/2025.05.08.652934

**Authors:** Pelin Icer Baykal, Mike Simonov, Dhrithi Deshpande, Ful Belin Korukoglu, Jaden Moore, Karishma Chhugani, Cecilia Liu, Varuni Sarwal, Neha Rajkumar, Mohammed Alser, Niko Beerenwinkel, Serghei Mangul

## Abstract

Genomic research relies on accurate and reproducible computational analyses of DNA sequencing data to draw reliable biological conclusions. Read mapping, the process of aligning reads to a reference genome, is central to many applications, including variant detection and comparative genomics. While several tools have been developed for this task, genomic reproducibility^1^, defined as the consistency of results across replicates, remains underexplored. Here, we address this question by introducing a methodology based on synthetic replicates of sequencing data, generated by perturbing the original reads through shuffling, reverse complementing, or combined shuffling and reverse complementing. Our approach is able to simulate variability observed across sequencing runs due to differences in library preparation techniques. We evaluated the reproducibility of eight alignment tools (BWA-MEM2^2^, Bowtie2^3^, HISAT2^4^, minimap2^5^, NextGenMap^6^, SNAP^7^, SMALT^7,8^ and Subread^9^) under these perturbations using whole-genome sequencing (WGS) data. Synthetic replicates were aligned and compared to the original sample to quantify discrepancies. Mapping accuracy changes ranged from 0.0001% to 4.4% for primary reads, which are alignments not marked as secondary, supplementary, or duplicates, and up to 12.2% for high-quality primary reads. For primary reads, the percentage of reads commonly mapped in both the original and synthetic replicate ranged from 91.66% to 100% relative to the total number of mapped reads in the original dataset. High-quality filtering improved consistency, though some tools still failed to recover more than 70% of the original alignments. Within the set of common reads, the incidence of inconsistent mappings was as high as 13.53% for primary reads and 6.73% for high-quality primary reads. Bowtie 2 was fully reproducible under the shuffling replicate, Subread was fully reproducible under the reverse-complement replicate, whereas NextGenMap exhibited only minor inconsistencies. By contrast, SNAP and minimap2 displayed the most significant variability under reverse complementing. We further demonstrate that alignment inconsistencies propagate to the downstream task of calling structural genomic variants. Using Manta for structural-variant (SV) calling, we observed that Bowtie 2, HISAT2, and minimap2 maintained perfect SV concordance between original and replicate alignments, whereas other tools exhibited lower concordance, with Subread scoring only 87% concordance. In conclusion, our comprehensive evaluation demonstrates that synthetic perturbations reveal critical differences in how alignment tools handle technical variability and how these differences propagate to downstream variant analyses, underscoring the necessity of incorporating reproducibility benchmarks into the selection and validation of read mappers to ensure robust and reliable genomic interpretations.

## Background

The process of aligning reads obtained from a sequencing experiment to a reference genome, known as read mapping, is fundamental in the field of genomics. Advances in high-throughput sequencing technologies have dramatically increased the volume and complexity of data generated^10^, placing significant emphasis on accurate and efficient read alignment to enable reliable downstream analyses, including variant detection, structural variant identification, and comparative genomic studies. Consequently, the accuracy and effectiveness of these alignment tools directly impact the validity of biological interpretations and subsequent conclusions derived from genomic data^11–13^. Given this critical role, rigorous evaluation of these tools is necessary to ensure consistency and reliability of genomic analyses.

Traditionally, benchmarking of alignment tools has emphasized metrics such as accuracy and computational efficiency, but less attention has been paid to genomic reproducibility—defined as the consistency of computational results with respect to technical variation that commonly occurs in sequencing experiments^1^. Interestingly, it has been observed that the computational tools and their configurations can have a more significant influence on genomic results than experimental factors like sequencing platform choice or library preparation methods^14^.

Despite the recognized importance of genomic reproducibility, systematically assessing the impact of subtle technical variations remains challenging. Although there are some efforts to generate technical replicates experimentally^14–16^ by sequencing the same biological sample multiple times through independent library preparation, this approach remains resource-intensive and often impractical, particularly in clinical research contexts or studies involving ethically sensitive samples, where repeated experimentation may be restricted due to ethical considerations or limited sample availability.

Existing evaluations of read alignment tools^11,17^ focus on accuracy and performance but rarely examine variations at different levels of alignment detail. Specifically, they do not distinguish clearly between global alignment (the exact genomic position where a read maps) and local alignment (the detailed matching of individual bases within the alignment). Evaluations typically concentrated on ambiguous reads (multi-mapped reads), where alignment algorithms may randomly assign reads to multiple equally plausible mapping locations, but they did not establish a systematic approach for assessing genomic reproducibility^18^. In some cases, tools report random alignments for the same read across multiple runs, confounding reproducibility assessments. Monitoring these discrepancies is crucial for understanding how technical perturbations can affect read alignments in ways that potentially impact downstream analyses.

To close this gap, we introduce a novel approach that employs three synthetic replicates: shuffling synthetic replicate, reverse complementing synthetic replicate, and combined perturbation synthetic replicate to systematically evaluate the genomic reproducibility of alignment tools. These synthetic replicates simulate realistic sources of technical variability that may naturally occur during sequencing experiments, such as changes in read order or orientation due to library preparation protocols or as a result of different sequencing runs^19,20^. While shuffling tests whether tools produce consistent results despite changes in read input order, reverse complementing introduces strand inversion, enabling us to probe how tools handle strand-specific alignment. This is particularly important because alignment tools treat strand information and strand bias, meaning preferential alignment of reads to one strand over another^21^, differently. Although other perturbation strategies such as bootstrapping or subsampling exist^1^, they disrupt the one-to-one correspondence between original and perturbed reads, making them unsuitable for pairwise reproducibility assessments in read alignment.

We assess genomic reproducibility across five WGS datasets of paired-end Illumina short reads and conduct a comprehensive benchmarking of eight non-stochastic read alignment tools: BWA-MEM2^2^, Bowtie2^3^, HISAT2^4^, minimap2^5^, NextGenMap^6^, SNAP^7^, SMALT^7,8^ and Subread^9^. By comparing alignments of synthetic replicates to their corresponding original datasets, our approach allows a comprehensive evaluation of reproducibility at both global (mapping locations) and local (edit distances) levels.

Reproducibility in read alignment is crucial, because alignment variability may impact downstream genomic analyses. For example, detecting structural variants (SVs) from DNA sequencing data relies heavily on precise read alignments. Minor inconsistencies in mapping locations or alignment qualities can lead to differences in SV calls. Evaluating how alignment reproducibility affects SV calling outcomes is therefore essential for fully understanding their reliability. We extend our reproducibility assessment framework to quantify how synthetic perturbations in read alignment propagate into downstream SV calling and find that the impact of shuffling on read alignment tool performance is also reflected in structural variant detection results, demonstrating the immediate impact of alignment reproducibility on SV detection using the shuffling synthetic replicate. Collectively, these findings underscore genomic reproducibility as a critical benchmark, complementing traditional performance metrics, and provide practical insights to guide researchers in selecting robust and reliable alignment tools for genomic analyses.

## Results

### Assessing genomic reproducibility of read alignment tools

We developed a methodology to evaluate genomic reproducibility of read alignment tools using whole-genome sequencing (WGS) synthetic replicates that simulate realistic technical variation. These synthetic replicates were generated from five paired-end Illumina datasets (Table S1) by applying three controlled perturbations to the original reads: shuffling synthetic replicate, reverse complementing synthetic replicate, and combined perturbation synthetic replicate (Fig. 1). Each perturbation represents a common source of technical variability encountered in sequencing workflows, such as differences in read ordering or strand orientation.

**Figure 1:**
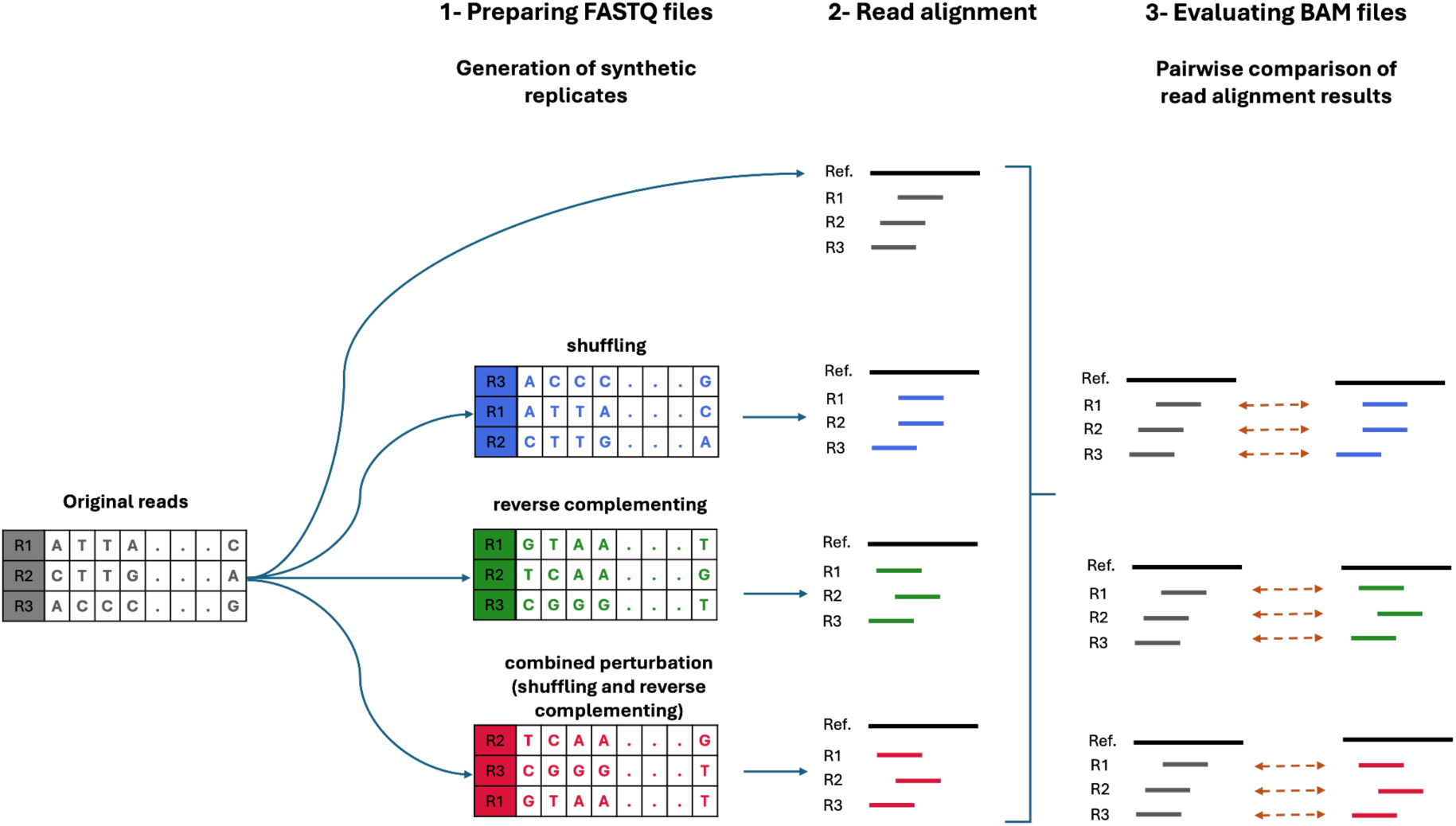
Workflow for assessing read alignment reproducibility. The process consists of three steps: (1) generating synthetic replicates from the original FASTQ data, (2) aligning both the original and synthetic replicate reads, and (3) performing pairwise comparisons of the alignments to evaluate reproducibility.

We systematically evaluated the reproducibility of eight widely used short-read alignment tools: BWA-MEM2^2^, Bowtie2^3^, HISAT2^4^, minimap2^5^, NextGenMap^6^, SNAP^7^, SMALT^7,8^ and Subread^9^ (Table S2) selected to reflect a broad range of algorithmic strategies, including Burrows-Wheeler Transform, hash-based methods, and seed-and-extend heuristics^21^. These tools were run with standardized parameters and deterministic configurations to eliminate any non-reproducibility stemming from tool-internal randomness (e.g., multithreading) (Table S3).

Since some reads can match equally well to multiple locations in the genome, tools often face challenges in determining a single “best” alignment. This can introduce ambiguity and make it harder to assess how consistently a tool behaves. To ensure alignment comparisons focused on reliable results, we restricted our analysis to primary alignments, and in a second tier, to high-quality primary reads defined as those aligned in proper pairs (i.e., both ends of a paired-end read mapping to the expected orientation and genomic distance) with maximum mapping quality scores (Table S4). This filtering reduces the influence of multi-mapped or ambiguous reads and ensures that reproducibility assessments reflect the most confident alignments. Across tools, the proportion of primary mapped reads among all reads was generally high (>90% for most tools), while the fraction of high-quality primary reads among primary reads varied more widely from ∼94% for HISAT2 to only ∼26.5% for Subread (Fig. S1).

We assessed reproducibility by comparing alignment outcomes between each original dataset and its corresponding synthetic replicates. We first evaluated the effect of perturbations on basic mapping behavior by calculating mapping rate differences, i.e., the change in the percentage of reads successfully aligned. We then examined consistency among commonly mapped reads, i.e., reads that appear in both original and replicate alignments, and quantified discrepancies in mapping location and edit distance. A read was considered inconsistent if either its genomic location or edit distance differed between the original and perturbed alignments. Although subtle alignment differences, such as variations in mismatch or gap placement within an alignment, could theoretically occur at identical genomic locations with the same edit distance, these cases were not separately considered. These inconsistencies were further classified into three types: consistent global – inconsistent local, where the read maps to the same position but with different alignment details; inconsistent global, where both the position and alignment details differ; and alternate location where the read maps to a completely different location (Fig. 2).

**Figure 2:**
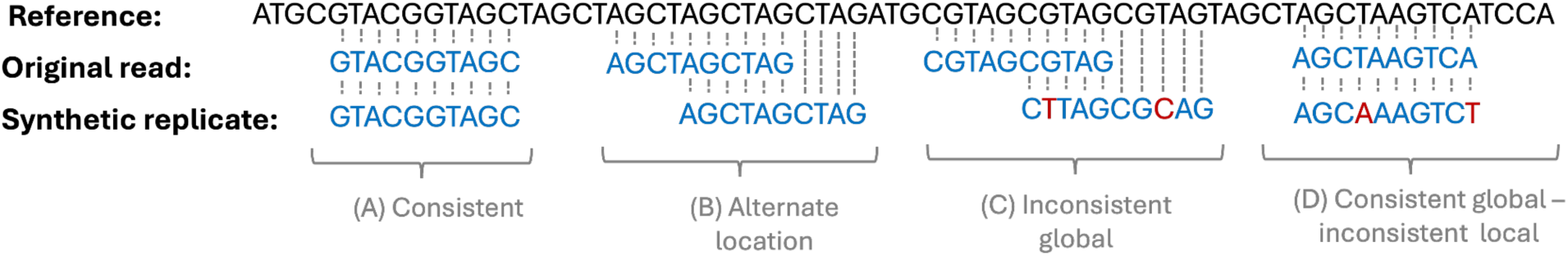
Classification scheme for read alignment inconsistencies. Reads are categorized as (A) Consistent: identical alignment between replicates, (B) Alternate location: mapped to a different but identical sequence, (C) Inconsistent global: mapped to different locations with sequence mismatches, and (D) Consistent global – inconsistent local: aligned to the same global region but with local sequence differences.

To summarize overall tool performance, we classified reproducibility outcomes based on two dimensions: commonality and consistency. Mapping commonality measures the proportion of reads mapped in both the original and replicate datasets, while consistency measures whether common reads align identically. We define full commonality as 100% of reads mapped in common, and partial common as less than 100%. Similarly, full consistency indicates 100% identical alignments among common reads, and partial consistency indicates non-zero discrepancies. These classifications are summarized in Table S5.

### The majority of the tools are able to maintain consistent mapping rates across replicates

To assess the effect of synthetic perturbations on read alignment rates, we compared the fraction of reads mapped between each original dataset and its corresponding synthetic replicates (Fig. S2).

For primary reads, shuffling caused minimal changes across all tools. For example, BWA-MEM2 showed a mean mapping-rate difference of 2.10 × 10⁻⁵, while Bowtie2, HISAT2, minimap2, and SNAP had differences of 0. Subread and NextGenMap showed slightly larger but still minor deviations (Fig. S2). High-quality primary reads exhibited similar stability, with only very small variations across tools.

In contrast, reverse complementing had a more noticeable impact. For primary reads, minimap2 and SNAP showed mean mapping-rate differences of 2.73 % and 4.42 %, respectively. Among high-quality reads, minimap2 was the most affected, with a difference of 12.23 %, whereas other tools remained below 5 % (Fig. S2).

Overall, most tools maintained stable mapping rates across replicates, except for minimap2 and SNAP under the reverse complementing synthetic replicate.

### Reproducibility of alignment outcomes under synthetic perturbations

To evaluate the reproducibility of alignments under synthetic perturbations, we quantified the percentage of common mapped reads, defined as reads successfully aligned in both the original and replicate datasets, relative to the number of reads mapped in the original (Fig. 3, Table S5). Among the reads that were mapped for both original data and synthetic replicate, we further assessed the proportion of these common reads that were inconsistently aligned, either due to differences in mapping location or edit distance (Fig. 4, Table S5).

**Figure 3:**
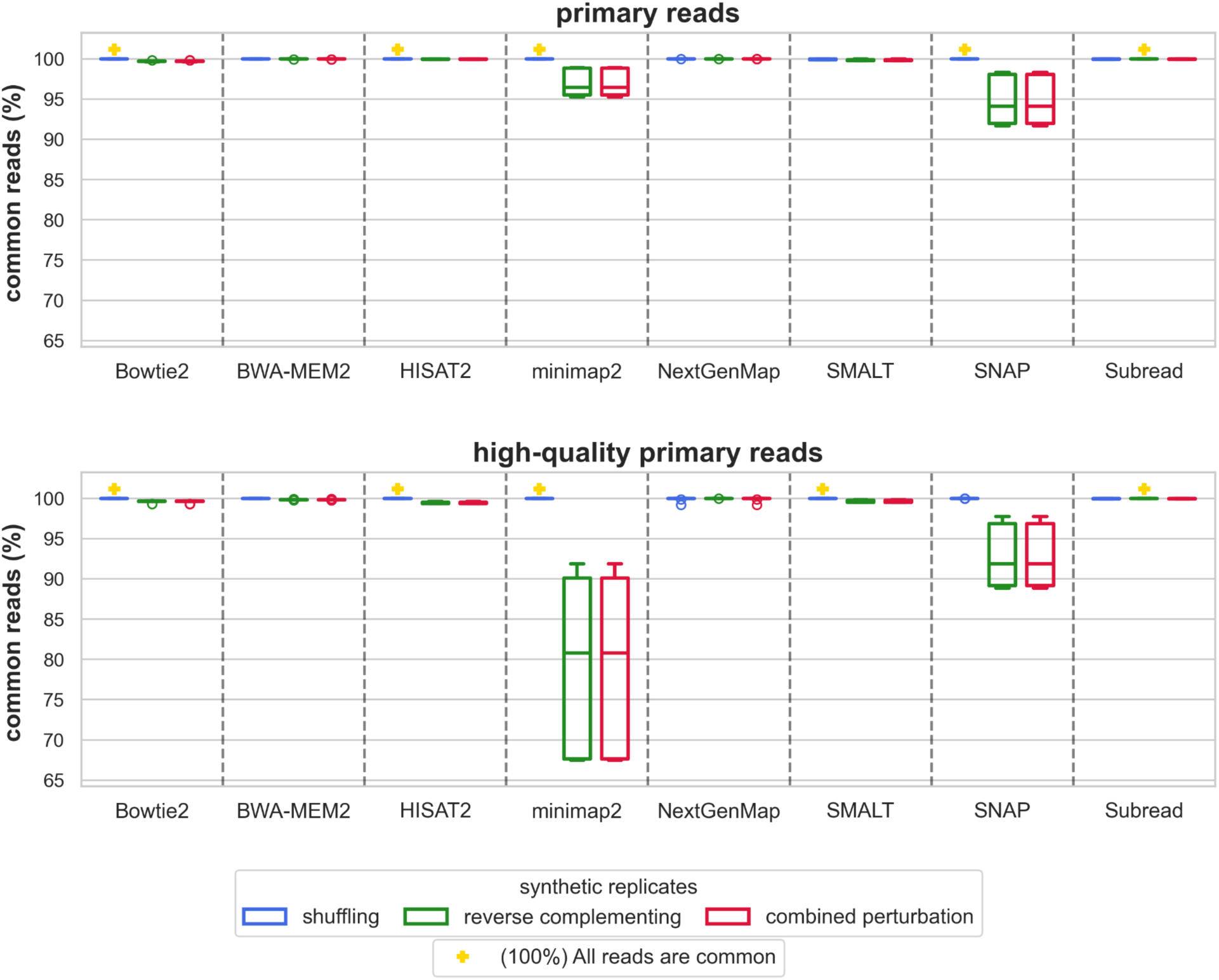
Percentage of common mapped primary reads between original sample and synthetic replicate. The top panel represents all primary reads, while the bottom panel focuses on high-quality primary reads only. Boxplots represent the distribution of common mapped reads across different alignment tools. Synthetic replicate types—shuffling (blue), reverse complement (green), and reverse complement with shuffling (red)—are compared. Yellow markers represent an average of 100% common mapped reads.

**Figure 4:**
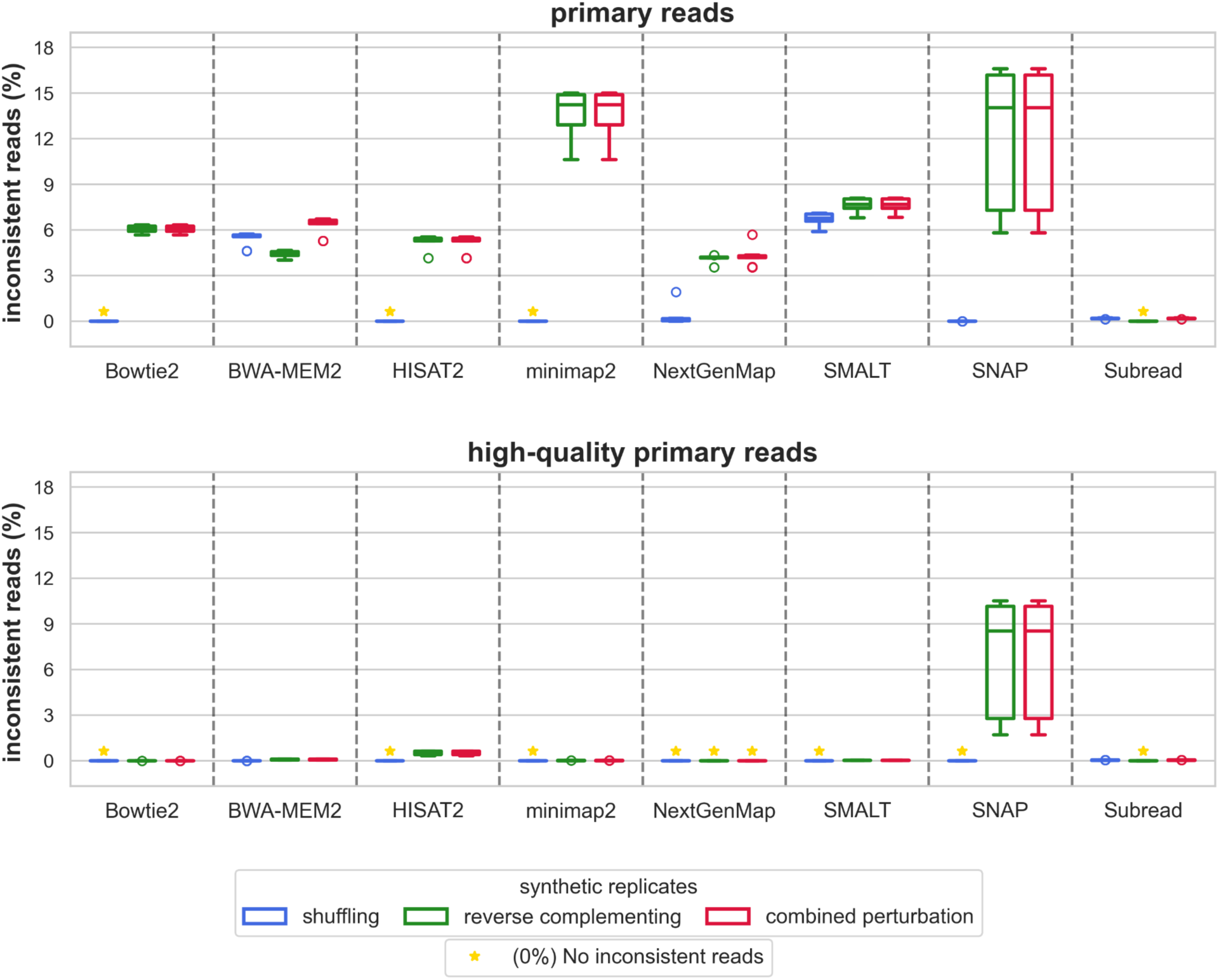
Percentage of inconsistent reads between original and synthetic replicate. The top panel shows results for all primary reads, while the bottom panel focuses on high-quality primary reads only. Boxplots represent the distribution of inconsistently mapped reads across different alignment tools. Synthetic replicate types—shuffling (blue), reverse complement (green), and reverse complement with shuffling (red)—are compared. Yellow markers represent an average of 0% inconsistent reads.

For primary reads under the shuffling synthetic replicate, HISAT2, minimap2 and Bowtie2 mapped 100 % of the original reads in common and showed 0 % inconsistencies, achieving full commonality and full consistency. SNAP also exhibited 100% commonality but with a marginal inconsistency rate (∼99.9% consistency). NextGenMap, SMALT, Subread, and BWA-MEM2 showed partial commonality (∼99.9 %) and inconsistency rates ranging from 0.18 % to 6.7 %. When high-quality reads were analyzed instead, reproducibility improved. HISAT2, minimap2, Bowtie2, and SMALT maintained full commonality and full consistency. SNAP and NextGenMap showed partial commonality (∼99.9 %) and full consistency. Subread and BWA-MEM2 continued to exhibit partial commonality and minor inconsistencies. Across these tools, excluding minimap2 and SNAP, the average proportion of common reads remained above 99%, highlighting that most reads are reliably mapped despite minor inconsistencies.

Reverse complementing introduced more pronounced variability. For primary reads, only Subread maintained full commonality and full consistency. All other tools, including HISAT2, minimap2, SNAP, SMALT, Bowtie2, NextGenMap, and BWA-MEM2, showed partial commonality (e.g., ∼94.8 % for SNAP and ∼96.9 % for minimap2) and inconsistency rates ranging from ∼4 % to 13.5 %. Among high-quality reads, Subread again achieved full commonality and full consistency. NextGenMap showed partial commonality (99.99 %) and full consistency. Other tools continued to exhibit variability. Minimap2 showed partial commonality (79.6 %) despite high consistency (99.99 %), while SNAP reached partial commonality (92.9 %) and partial consistency (93.3 %). Bowtie2, SMALT, HISAT2, and BWA-MEM2 similarly retained partial commonality and partial consistency due to non-zero inconsistency rates.

These results demonstrate that while several tools maintain high reproducibility under the shuffling synthetic replicate, the reverse complementing synthetic replicate introduces greater alignment variability—particularly for tools such as minimap2 and SNAP that exhibit sensitivity to strand orientation. High-quality filtering generally improves consistency but does not always restore full commonality, highlighting the importance of evaluating both mapping agreement and alignment stability when assessing genomic reproducibility.

To supplement these results, we visualized the relative proportions of common, original-only, and replicate-only mapped reads using a union-based breakdown (Fig. S4). While most tools showed high commonality across replicates, tools such as SNAP and minimap2 exhibited larger fractions of original-only reads, particularly under reverse complementation and combined perturbations. This visualization offers an intuitive perspective on mapping asymmetries across tools and conditions, complementing the primary reproducibility metrics.

### Effect of combined perturbations on alignment reproducibility

To quantify how the combined perturbation differs from each individual perturbation (shuffling synthetic replicate and reverse complementing synthetic replicate), we calculated the Jaccard index between those from the shuffling synthetic replicate and the reverse complementing synthetic replicate (Fig. 5). A lower Jaccard index suggests that combined perturbation introduced new inconsistencies not seen with either perturbation alone, while a higher index indicates similarity in behavior across conditions.

**Figure 5:**
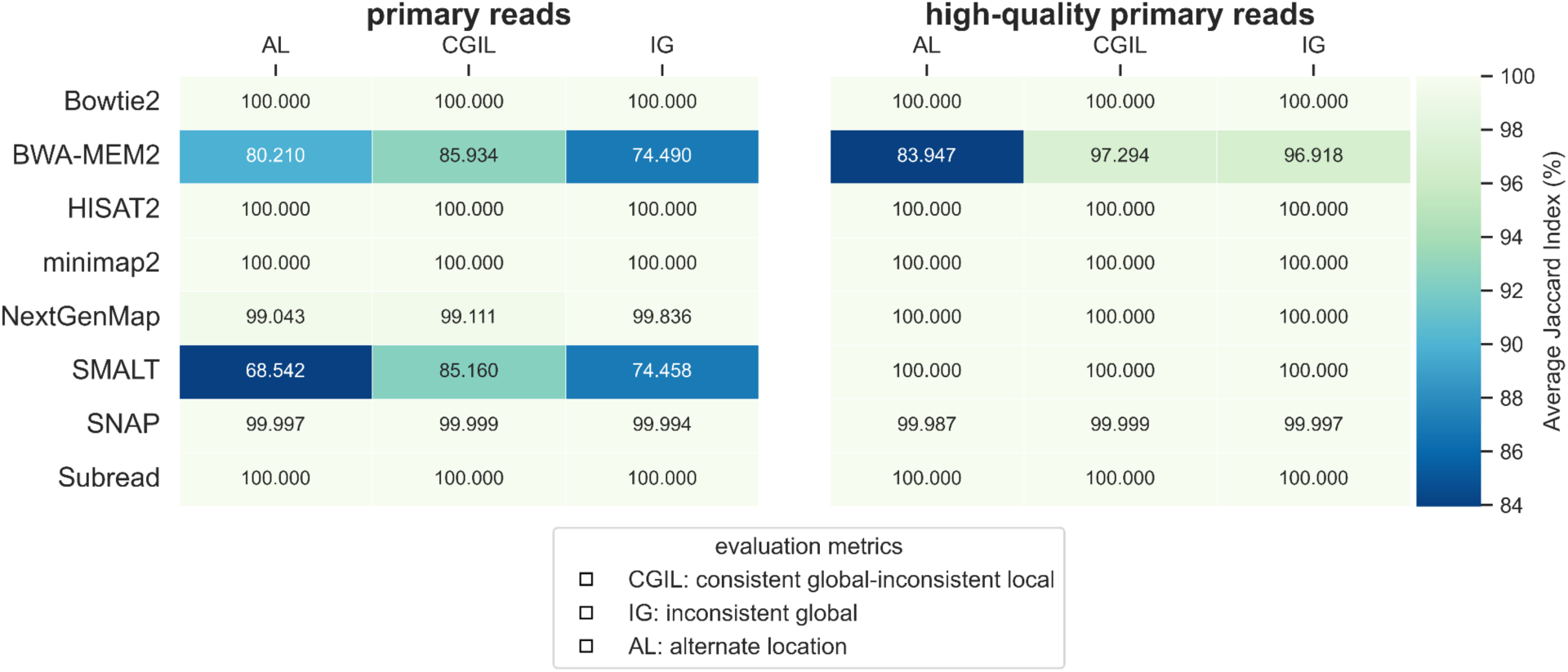
Jaccard similarity comparing the set of inconsistent reads under the combined perturbation (shuffling + reverse complementing) to each individual perturbation. The left panel represents all primary reads, while the right panel focuses on high-quality reads. A Jaccard index below 1 indicates that combining shuffling and reverse complementing introduces additional mapping variability.

For primary reads, BWA-MEM2 and SMALT showed similar variability under the combined perturbation, with Jaccard indices of 80.21%, 85.93%, and 74.49% for BWA-MEM2, and 68.54%, 85.16%, and 74.46% for SMALT corresponding to alternate location, consistent global–inconsistent local, and inconsistent global categories, respectively. In contrast, NextGenMap and SNAP had values approaching 99%, indicating high consistency between the combined perturbation synthetic replicate and the individual perturbation synthetic replicates. Other tools remained at 100%, suggesting no additional differences under the combined perturbation (Fig. 5).

For high-quality reads, Jaccard similarity values were generally higher than those observed for primary reads, indicating increased consistency between the sets of inconsistent reads found in the combined perturbation synthetic replicate and those found in individual perturbations (Fig. 5). However, SNAP continued to show some residual variability. These results indicate that for certain tools, combined perturbation synthetic replicate introduces more inconsistencies than either shuffling or reverse complementing alone, emphasizing the need to assess reproducibility under multiple forms of technical variation.

To complement the original-relative metrics used throughout our reproducibility assessment, we additionally summarized the read alignment results based on the union of mappings across the original and replicate datasets (Fig. S4). Specifically, for each tool and synthetic replicate type, we calculated the proportion of reads that were mapped in both datasets (common reads), only in the original dataset (original-only reads), or only in the replicate dataset (replicate-only reads). This union-based breakdown provides a more detailed view of reproducibility failures by distinguishing between lost alignments (reads missing in the replicate) and new alignments (reads appearing only in the replicate). While most tools exhibited minimal imbalance, certain cases revealed pronounced asymmetries. For example, minimap2 under reverse complementation failed to recover over 20% of high-quality alignments originally present, while producing few new replicate-only mappings. In contrast, Bowtie2 and SMALT showed more symmetric discrepancies, with both original- and replicate-only mappings around 0.4–0.5%. These patterns help clarify the directionality of alignment mismatches and highlight tool-specific sensitivities to input perturbations.

### Alignment tools exhibit discrepancies in both the overall mapping location and fine-scale alignment

To gain further insight into the nature of reproducibility failures, we classified commonly mapped reads according to the three evaluation metrics (defined in the section ‘Mapping rate and reproducibility evaluation metrics’) (Fig. 2) and visualized their proportions (Fig. 6). Among primary reads, alternate location mismatches were the most frequent inconsistency across most tools, indicating that shifts in mapping position were the main driver of failures. Subread showed a comparatively higher fraction of consistent global inconsistent local reads, while minimap2 exhibited a more even split between inconsistent global and alternate location reads (Fig. 6).

**Figure 6:**
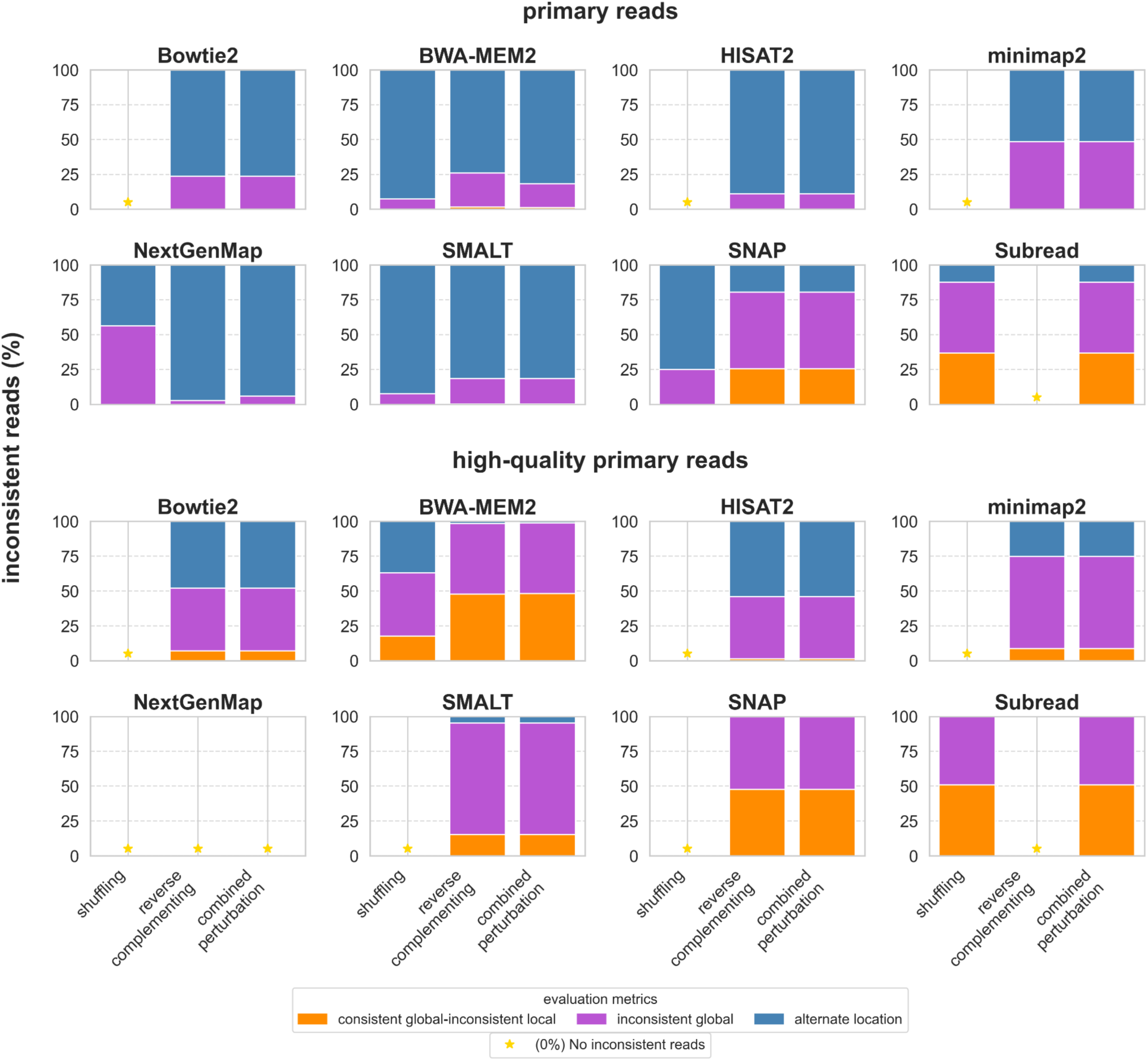
Breakdown of inconsistent reads across alignment tools using three evaluation metrics. Stacked bar plots show the proportion of inconsistent reads classified as Consistent Global – Inconsistent Local (CGIL, orange), Inconsistent Global (IG, blue), and Alternate Location (AL, green). The top panel represents all primary reads, while the bottom panel focuses on high-quality primary reads only. Yellow markers represent 0% inconsistent reads.

In the high-quality subset, the total number of inconsistencies decreased for all tools, but the relative breakdown between consistent global inconsistent local, inconsistent global, and alternate location mismatches remained tool specific. Bowtie2 was still dominated by alternate location mismatches, while BWA-MEM2 maintained a balanced mix of inconsistent global and consistent global inconsistent local mismatches. These patterns illustrate that different aligners vary not only in how often they fail to reproduce mappings but also in the types of failures they incur, emphasizing the importance of considering both global and local alignment variability when selecting an alignment tool.

### Backward reads are more sensitive to perturbations

Strand orientation can influence how reads are aligned, so we specifically assessed whether reads originating from the forward strand versus the reverse strand exhibit different inconsistency patterns under synthetic perturbations. Reverse complementing applies a reverse-complement transformation to each read, enabling evaluation of whether alignment tools demonstrate differential sensitivity to forward- and reverse-oriented reads (Fig. S3).

Overall, the high-quality subset showed reduced discrepancies across tools, supporting that stringent filtering improves alignment stability. However, across all tools, backward reads consistently exhibited greater sensitivity to perturbations than forward reads. This indicates that alignments for reverse-oriented reads are more prone to shifts in mapped location or changes in edit distance.

For primary reads, Bowtie2 was particularly sensitive to strand inversion, with mean locational differences reaching 33.7 Mb for backward reads compared to 31.7 Mb for forward reads. In contrast, SNAP maintained perfect locational reproducibility with the shuffling synthetic replicate. Subread demonstrated the most stable performance, with differences in mapped genomic positions remaining below 5 Mb across all perturbations.

When considering high-quality reads, locational discrepancies decreased for nearly all tools. However, backward reads remained more variable than forward reads. BWA-MEM2 and minimap2 showed increased shifts primarily in backward reads, whereas SMALT exhibited more variability in forward reads. Differences in edit distance followed a similar pattern: Bowtie2, minimap2, and SNAP showed large outlier discrepancies for primary reads, which were substantially reduced after high-quality filtering, indicating improved alignment fidelity under stringent conditions. Notably, all inconsistencies were eliminated with NextGenMap in the high-quality subset, resulting in no remaining mismatches to report.

### Downstream impact of alignment reproducibility on structural variant detection

To assess how non-reproducible read alignments affect downstream analyses, we examined the consistency of structural variant (SV) detection using the Manta SV caller. Specifically, we compared SVs identified from alignments of the original datasets to those detected from their shuffling synthetic replicate alignments.

To quantify reproducibility, we computed two concordance metrics and reported them as percentages. Replicate concordance (Rep→Orig) represents the percentage of SVs found in the replicate alignment that are also detected in the original alignment. Original Concordance (Orig→Rep) reflects the percentage of SVs found in the original alignment that are also recovered in the replicate. Higher values in both measures indicate greater reproducibility of SV detection under alignment perturbations (Fig. 7).

**Figure 7:**
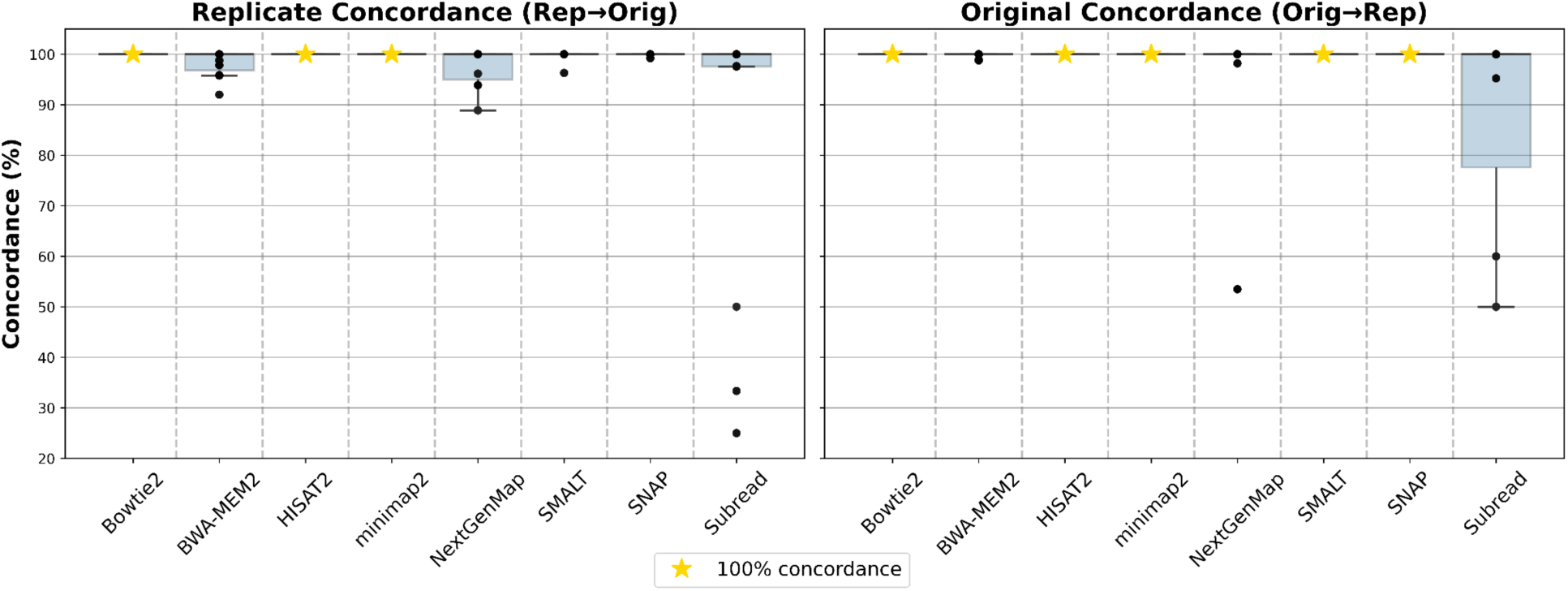
Reproducibility of structural variant detection across alignment tools. Replicate Concordance (Rep→Orig) and Original Concordance (Orig→Rep) percentages were calculated by comparing structural variants and indels detected from original alignments against those identified in shuffled synthetic replicates. All results were generated using the SV caller Manta. Higher concordance percentages indicate greater reproducibility in downstream variant detection under technical perturbations.

Bowtie2, HISAT2, and minimap2, which had previously shown full overlap and full consistency at the alignment level, maintained perfect reproducibility in downstream SV detection, achieving 100% concordance in both directions across all datasets. This outcome confirms that Manta behaves deterministically with identical alignment inputs and demonstrates that reproducible alignments translate into stable downstream variant calls.

Other tools did exhibit reductions in SV concordance. BWA-MEM2 achieved Replicate and Original Concordance percentages of 98% and 99%, respectively, differing from the original in a few missed or additional SVs. NextGenMap displayed even lower average scores (∼97%), with broader variability in Original Concordance, indicating increased omissions in replicatese- based SV detection. SNAP and SMALT were similarly inconsistent (mean replicate concordance: 99%; original concordance: 100%), and a small number of SVs detected only in the replicates reduced replicate concordance.

Subread exhibited the lowest reproducibility, with average concordance percentages around 87% in both directions. In some datasets, replicate concordance dropped as low as 25%, indicating that a large fraction of SVs detected in the replicates were not found in the original. Similarly, original concordance fell to 50%, meaning that half of the SVs in the original dataset were not recovered in the replicate. These pronounced discrepancies underscore how even minor differences in alignment can propagate and significantly impact downstream variant detection, reinforcing the importance of alignment reproducibility when selecting tools for genomic analyses.

## Discussion

Reproducibility in genomics is essential for ensuring that findings remain reliable even when technical variations occur during sequencing or data processing. In this study, we introduced three synthetic replicate types—shuffling, reverse complementing, and combined perturbation—to systematically evaluate the reproducibility of read alignment tools. This framework enables precise simulation of specified technical variations, such as changes in read order or strand orientation, and thus permits assessment of whether alignment tools produce consistent results under these defined perturbations. High reproducibility is characterized by minimal differences in key mapping outcomes, including mapping locations and edit distances, across synthetic replicate types.

Our results demonstrate that reproducibility varies substantially with the type of perturbation applied. For example, Bowtie2 maintained perfect reproducibility under the shuffling synthetic replicate but exhibited reduced consistency under the reverse complementing synthetic replicate, indicating sensitivity to strand inversion. Conversely, Subread remained fully reproducible by achieving full commonality and full consistency under the reverse complementing synthetic replicate but showed increased variability under shuffling. SMALT, HISAT2, and NextGenMap achieved high commonality and consistency across both perturbations, particularly after restricting to high-quality reads. These observations highlight the necessity of considering robustness to technical fluctuations alongside traditional accuracy and performance metrics when selecting an alignment tool.

An important distinction observed in our analysis is the difference between the proportion of inconsistent reads and the magnitude of their locational or alignment shifts. Some aligners, such as Bowtie2, maintained high consistency overall but experienced larger locational deviations for the few reads affected. Conversely, aligners such as SMALT and minimap2 exhibited smaller locational differences across a larger proportion of reads, distributing their variability more broadly. Thus, reproducibility assessments must account not only for the frequency of inconsistencies but also for the extent of their impact on alignment results.

Under the combined perturbation, NextGenMap and SNAP continued to demonstrate near-perfect commonality and consistency, whereas BWA-MEM2 and SMALT incurred new misalignments that were not observed under the shuffling or reverse complementing synthetic replicate alone. This behavior suggests that the alignment strategies of BWA-MEM2 and SMALT depend on assumptions about read order or strand orientation, and it illustrates why assessing multiple, distinct perturbations is essential for a comprehensive evaluation of tool robustness.

Orientation-specific analysis revealed that reverse-oriented reads consistently exhibited higher inconsistency rates than forward-oriented reads even after high-quality filtering. For instance, Bowtie2’s reverse reads were misaligned more frequently than its forward reads, whereas Subread treated both orientations with similar stability. These strand-dependent biases suggest that alignment algorithms may handle strands differently and that future development should aim to minimize such asymmetries.

In addition to our primary commonality and consistency metrics, we assessed mapping asymmetry through a union-based breakdown. We evaluated the proportion of reads that were aligned exclusively in the original dataset, exclusively in the synthetic replicate, or in both (Fig. S4). This breakdown provides a clearer view of how mapping discrepancies arise, revealing that for some tools, particularly minimap2 and SNAP, most mismatches stem from reads present only in the original alignment, a pattern that is especially pronounced in the high-quality subset.

Further analysis of downstream structural variant detection using Manta demonstrated that alignment reproducibility directly impacts variant calls. Bowtie2, HISAT2, and minimap2 maintained 100% replicate and original concordance, confirming that reproducible alignments yield stable variant detection. BWA-MEM2 and NextGenMap exhibited minor reductions to 98–99% concordance, whereas Subread’s concordance dropped to 87% in both directions, illustrating how alignment inconsistencies even at low levels can propagate into downstream analyses.

While synthetic replicates offer a controlled method for evaluating reproducibility, they have inherent limitations. Although effective in simulating read-order and strand-orientation variations, they do not fully capture the complexity of real-world sequencing variability, such as library preparation biases, sequencing chemistry artifacts, or instrument-specific errors. Additionally, our study focused on short paired-end Illumina reads; it remains unclear whether similar reproducibility patterns will persist in long-read sequencing technologies, which present distinct alignment challenges^22–24^. Extending this reproducibility framework to systematically benchmark multiple structural variant callers using synthetic perturbations, alongside incorporating more complex scenarios such as multi-mapping and ambiguous alignments, would provide deeper insights into genomic reproducibility.

In conclusion, our study demonstrates that controlled perturbations can reveal meaningful differences in how alignment tools handle data variability and how these differences affect downstream analyses. We advocate adopting reproducibility as a fundamental benchmark in genomic studies and recommend systematically evaluating both alignment tools and downstream analysis workflows under controlled perturbations. Tool developers should incorporate reproducibility metrics into their validation processes, thereby ensuring that genomic analyses remain robust and reliable under realistic sequencing conditions.

## Methods

### Choice of dataset and synthetic replicate design

To conduct our genomic reproducibility assessment of read alignment tools, we randomly selected five paired-end WGS Illumina short read datasets, each with a read length of 108 bp, from the 1000 Genomes Project (Table S1). These datasets provide high-quality, widely used genomic data suitable for benchmarking reproducibility.

We generated three types of synthetic replicates by random shuffling of reads (shuffling synthetic replicate), reverse complementing reads (reverse complementing synthetic replicate), and a combined perturbation (combined perturbation synthetic replicat) to emulate specific aspects of technical variation that have been shown experimentally to drive variability in sequencing analyses. In a multi–center study of structural variant calling, mapping methods accounted for the largest share of variability, followed by differences between sequencing centers and technical replicates; notably, calls supported by only one center or one replicate still overlapped the long-read SV call set at rates of 47.02 % and 45.44 %, respectively^22^.

Shuffling reflects the arbitrary order in which reads are captured and stored during sequencing and downstream processing. For example, during DNA hybridization on flow cells, single-stranded DNA fragments randomly bind to complementary oligonucleotides, resulting in spatially random clusters^18^. Moreover, in multiplexed sequencing, multiple samples are pooled together and sequenced simultaneously, and after demultiplexing, the reads from each sample are interleaved in an unpredictable order. This deliberate permutation preserves sequence content and quality scores but tests whether aligners produce consistent results under altered input order.

Reverse complementing simulates strand inversion by flipping both the bases and their quality scores, thereby mimicking scenarios in which reads originate from or are interpreted on the opposite strand. In real sequencing, such strand-bias issues can affect how tools handle alignments, making it important to assess reproducibility when reads are reversed. Finally, combined perturbation, combines shuffling and reverse complementing, evaluating each tool’s stability under a more complex scenario of altered ordering and orientation. While these modifications do not mirror every possible biological or technical event exactly, they capture typical ways in which data may vary and provide a controlled framework for analyzing how robustly read alignment tools handle such fluctuations.

### Read alignment tool selection and configuration

We selected eight state-of-the-art WGS read alignment tools: BWA-MEM2^2^, Bowtie2^3^, HISAT2^4^, minimap2^5^, NextGenMap^6^, SNAP^7^, SMALT^7,8^ and Subread^9^, available via the Bioconda channel^23^ These tools were selected based on their popularity in the genomics community, algorithmic diversity, and continued relevance in recent publications. All support DNA short-read alignment and are commonly employed in whole genome sequencing (WGS) pipelines. The selected aligners represent a range of mapping strategies such as Burrows-Wheeler Transform, hash-based indexing, and seed-and-extend heuristics.

We configured each tool with consistent parameters and, where necessary, adjusted settings to eliminate sources of non-determinism such as multithreading. Some aligners produce variable results across runs when executed with multiple threads; to avoid this, tools were executed in single-threaded mode where applicable (Table S3). Notably, minimap2 was used with its default three-thread configuration as it produced deterministic results under our settings. All tools aligned to the GRCh38 human genome assembly obtained from NCBI.

To accommodate strand-specific changes introduced by reverse complementing and combined perturbation, we adjusted tool-specific flags where applicable. Bowtie2, HISAT2, and Subread were run with the --rf flag, while SMALT was configured with the -l mp option. These options ensured proper interpretation of strand-specific data. BWA-MEM2, SNAP, minimap2, and NextGenMap do not have documented flags for reverse-complement handling and were run according to default recommendations.

### Selection of reads for analysis

Multi-mapping, where a read aligns to multiple genomic locations, poses a significant challenge to reproducibility, as it introduces ambiguity in determining the correct alignment. To mitigate this and ensure a fair and reliable comparison of alignment outcomes, we filtered the BAM files to retain only primary alignments and high-quality reads. Primary reads were defined as those not flagged as supplementary, secondary, or duplicate. We further defined high-quality reads as primary alignments that were mapped as proper pairs and assigned the maximum mapping quality score (MAPQ) by each alignment tool (Table S4). This filtering allowed us to restrict our reproducibility assessments to the most confident and unambiguous alignments.

### Mapping rate and reproducibility evaluation metrics

As a preliminary step to evaluating reproducibility, we assessed the effect of synthetic perturbations on basic alignment behavior. For each dataset and alignment tool, we computed the mapping rate difference between the original and its corresponding synthetic replicate. Mapping rate was defined as the percentage of input reads successfully aligned by the tool. The mapping rate difference was calculated as the mapping rate of the original dataset minus that of the synthetic replicate. Positive values indicate that fewer reads were aligned after perturbation, while negative values indicate more reads were aligned. This analysis was conducted separately for both primary and high-quality read sets.

To assess the genomic reproducibility of read alignment tools, we compared the consistency of read alignment outcomes between synthetic replicates and the original data. We first defined common mapped reads as those successfully aligned in both the original and synthetic replicate datasets, calculated as a percentage relative to the number of reads mapped in the original dataset. Inconsistent reads were defined as a subset of these common mapped reads whose alignment characteristics differed across datasets. We then defined specific categories of alignment differences (Fig. 1) to classify variations in mapping locations and edit distances. We classify a read’s alignment as inconsistent when its mapping location or edit distance differs between the original dataset and its synthetic replicate. In other words, if a read is mapped to a different genomic location or displays a different edit distance in the synthetic replicate compared to the original, its mapping outcome is considered inconsistent. This definition captures discrepancies both in the overall mapping location (global inconsistencies) and in the fine-scale alignment details (local inconsistencies). For paired-end data, consistency requires that both ends of a read pair are mapped to the same location with identical edit distances in both datasets; if either deviates, the pair is regarded as inconsistent. This systematic categorization allows us to assess the reproducibility of alignment outcomes across synthetic replicates and the original data.

To further refine our analysis of inconsistent read alignments, we developed a set of evaluation metrics to classify discrepancies between the original dataset and its corresponding synthetic replicates (Fig. 2). First, we identified the subset of reads that were successfully mapped on both the original and synthetic replicate alignments. For each commonly mapped read, we performed a pairwise comparison of its reported mapping location and edit distance. A read is classified as exhibiting ‘consistent global - inconsistent local’ alignment if it is mapped to the same genomic location in both datasets but displays a different edit distance. This indicates variability in fine-scale alignment details despite a stable overall locus. Conversely, if a read shows differences in both its mapping location and edit distance between the original and synthetic replicate alignments, it is classified as ‘inconsistent global’ (IG), reflecting discrepancies in both its overall placement and local alignment fidelity. Finally, if a read is assigned an alternative location in the synthetic replicate alignment relative to the original dataset, it is designated as an ‘alternate location’ (AL) read. This systematic, pairwise comparison of commonly mapped reads provides a precise and controlled framework for assessing the genomic reproducibility of read alignment outcomes at both global and local levels.

To provide an additional summary of mapping behavior, we calculated the percentage of reads that were mapped only in the original dataset, only in the replicate, or in both (common), relative to the union of all mapped reads across both datasets. This union-based breakdown offers a standardized, mutually exclusive partitioning of mapped read outcomes, enabling supplementary insights into directional differences in mapping behavior between the original and perturbed datasets (Fig. S4).

### Snakemake workflow

For our analysis, we developed a Snakemake workflow^24^, designed to create synthetic replicates and assess genomic reproducibility from WGS data (Fig. 1). This comprehensive workflow is designed to accommodate both single-end and paired-end data inputs in FASTQ format.

The workflow begins with the data preparation stage, where the input data format is standardized and FASTQ files are validated to ensure integrity. The validation checks whether the data corresponds to paired-end or single-end sequencing. Additionally, during this stage, it enumerates the total number of reads from the input FASTQ file and records these counts into a CSV file for subsequent analysis.

The alignment stages use eight pre-selected tools (Table S2) to map both original and synthetic reads to the reference genome. The parameters and settings for these tools were chosen based on their user manuals and refined through preliminary tests to ensure deterministic outcomes (Table S3). Specifically, we applied fixed seed values (where supported) and configured each tool with recommended settings for short-read alignment, and when necessary with additional flags to correctly handle strand-specific perturbations (reverse complementing and combined perturbation). These choices ensure that any observed differences in alignment results can be attributed to the synthetic replicate perturbations rather than to inherent variability in the tools.

The workflow parses each BAM file to extract key alignment attributes such as read identifiers, mapping locations, mapping quality scores, and edit distances, and stores these data in a structured CSV format. These CSV files form the basis for a read-by-read comparison between these data in a structured CSV format. For every read that is present in both datasets, the workflow evaluates discrepancies in its mapping location and edit distance according to the predefined categories (consistent global - inconsistent local, inconsistent global, and alternate location). The results of these pairwise comparisons are then aggregated into summary statistics that quantify overall genomic reproducibility of alignment tools, including metrics such as the percentage of inconsistent mappings and the average magnitude of differences in mapping characteristics. The final output is a comprehensible CSV file that details these comparative metrics, thereby enabling reproducibility and independent verification of the analysis.

The workflow offers high customizability through a configuration file that permits modifications of key parameters, such as the number of synthetic replicates and the choice of alignment tools. By default, the workflow is set to compare eight state-of-the-art alignment tools (Table S2). Each tool is governed by its own rule, written in separate Snakemake files (.smk files), enhancing the workflow’s modularity. This setup not only facilitates the maintenance and updating of individual alignment processes but also allows users to expand the comparison by adding more alignment tools. Users can integrate additional tools by providing the associated rules in separate Snakemake files.

### Analysis of the structural variant calling

To investigate how alignment reproducibility affects downstream analyses, we performed structural variant (SV) calling using the Manta tool. BAM files from original and shuffling synthetic replicate alignments were used as input. We restricted this analysis to shuffling synthetic replicates, since reverse complementing and combined perturbation involve strand perturbations that are incompatible with the assumptions of standard SV callers.

Following SV detection, VCF files were compared between original and replicate datasets. We quantified reproducibility using two concordance metrics and reported them as percentages. Replicate concordance (Rep→Orig) represents the percentage of SVs found in the replicate alignment that are also detected in the original alignment. Original concordance (Orig→Rep) reflects the percentage of SVs found in the original alignment that are also recovered in the replicate. These metrics provided a measure of how reliably each alignment tool preserved downstream variant detection outcomes under synthetic perturbations.

## Availability of data and materials

Code: https://github.com/cbg-ethz/GenoRepro/ Data: From 1000 Genomes Project: Access IDs: ERR009308, ERR009309, ERR009332, ERR009345, ERR013013. https://www.internationalgenome.org/

## Acknowledgments

We thank Aditya Sarkar for his valuable assistance with data analysis.

## Author contributions

Pelin Icer Baykal (PIB): Software; Formal analysis; Data curation; Visualization; Writing – original draft.

NB and SM: Supervision; Writing – review & editing. (Equal contribution) MS: Software; Investigation; Data curation; Visualization

MA: Supervision; Writing – review & editing. FBK, DD, NR, KC, CL: Investigation.

JM: Data curation; investigation VS: Resources.

## Supplementary Figures

**Figure S1:**
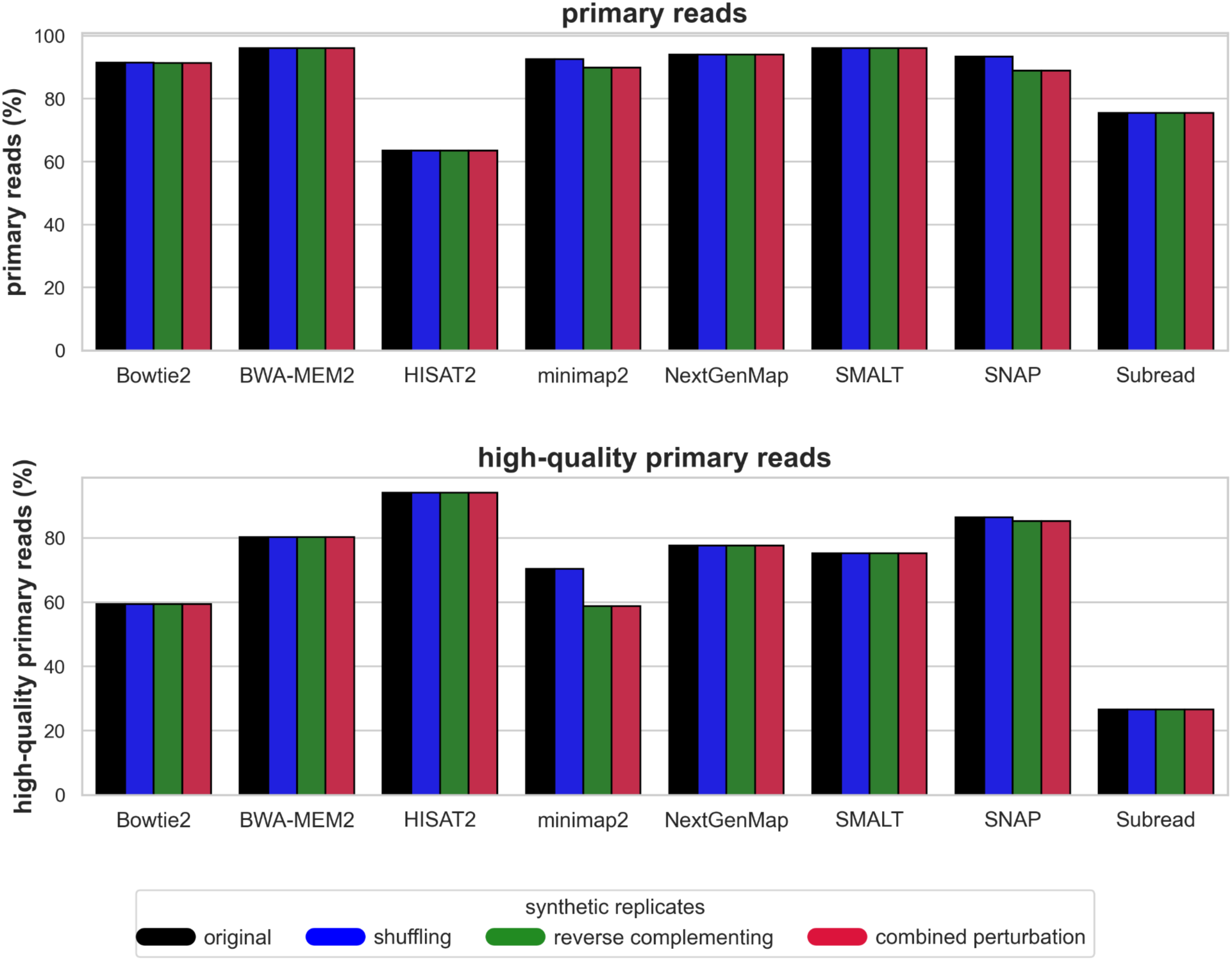
Proportion of mapped primary reads over all reads and proportion of mapped high-quality reads over all mapped primary reads. Bar plots representing the percentage of mapped primary reads among all reads and the percentage of mapped high-quality reads among mapped primary reads for each tool and its synthetic replicates.

**Figure S2:**
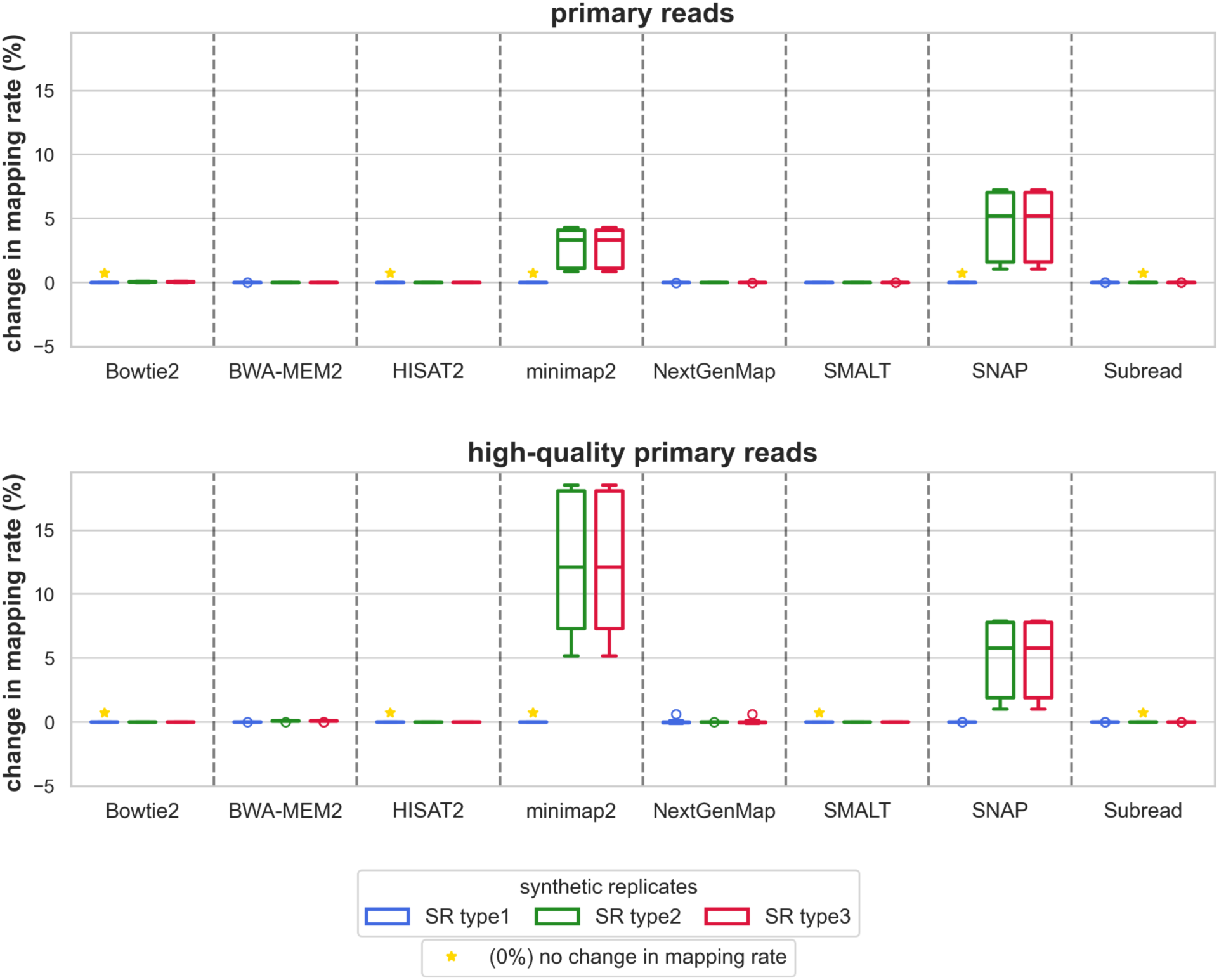
Mapping rate differences between original and perturbed datasets. Boxplots representing mapping rate difference of mapped reads between original dataset and its synthetic replicate dataset.

**Figure S3:**
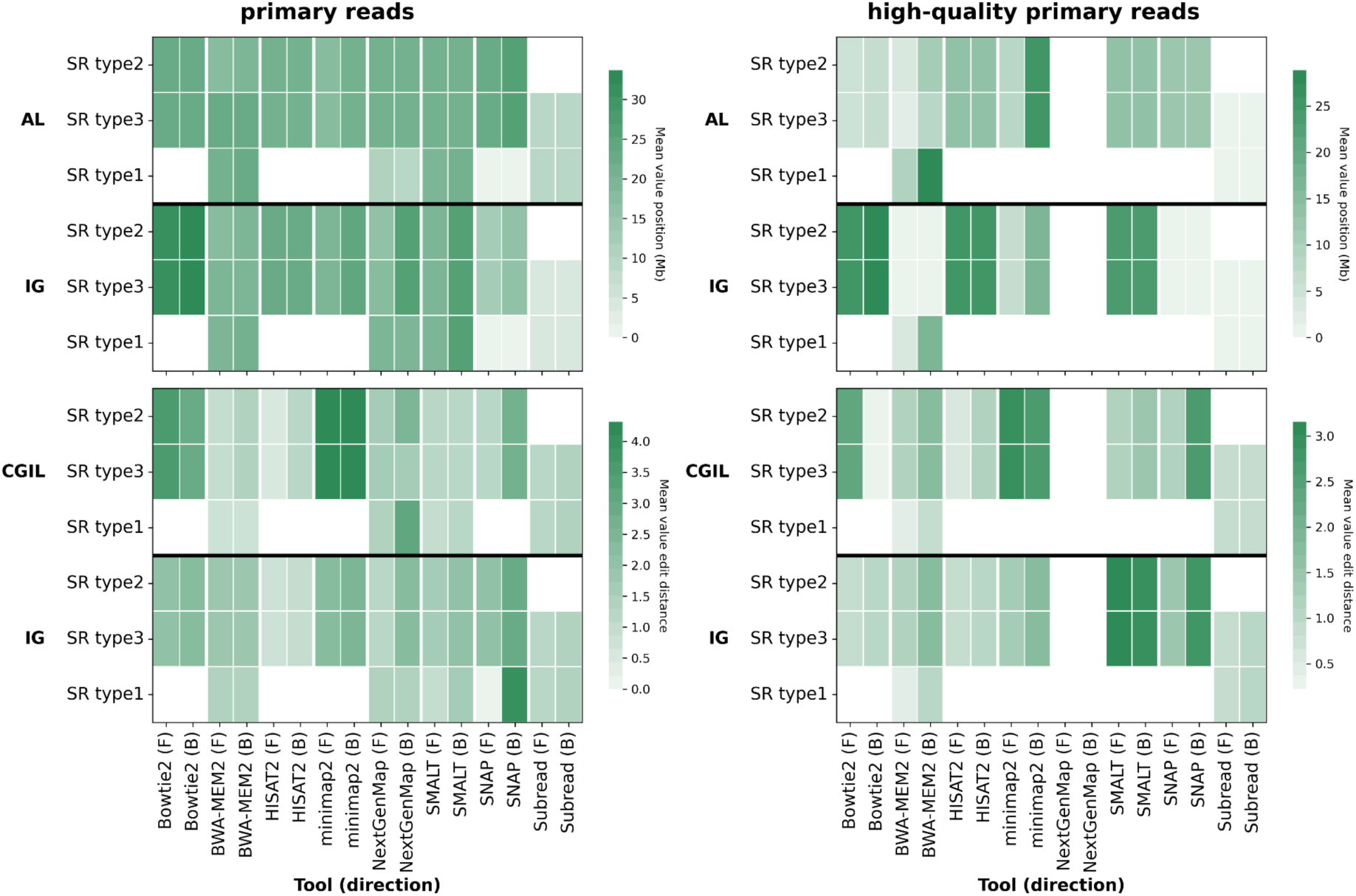
Locational and edit distance discrepancies by read orientation. Heatmaps display mapping location differences (top) and edit distance differences (bottom) across synthetic replicate types and aligners. The left panel shows primary reads, the right panel shows high-quality reads. F = forward reads, B = backward reads. Blank cells indicate perfect reproducibility. Blank cells indicate cases where all reads were identical for the corresponding alignment tool and synthetic replicate combination.

**Figure S4:**
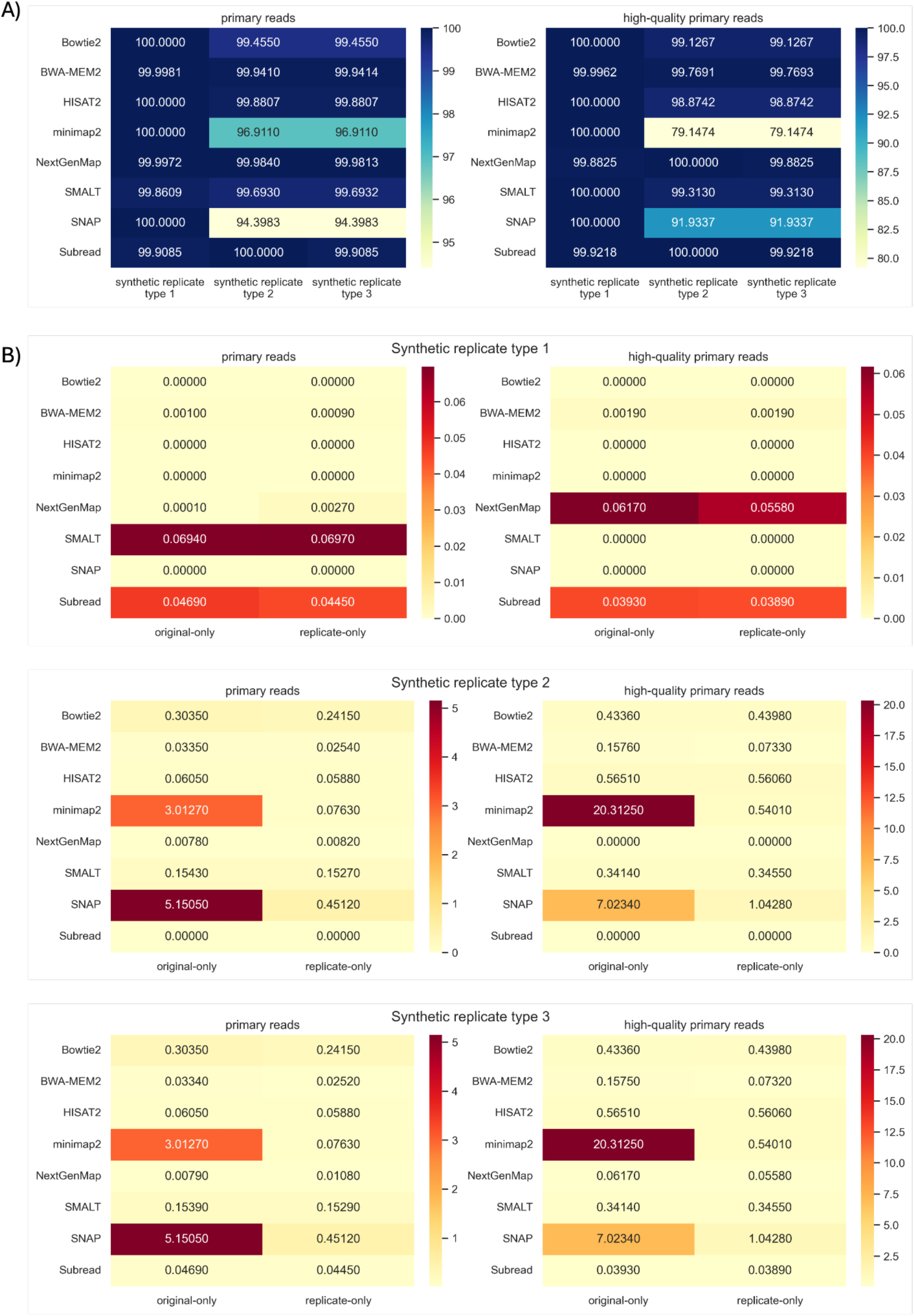
Union-based breakdown of read mapping overlap across synthetic perturbations. (A) Heatmaps of the mean percentage of reads commonly mapped in both the original dataset and its synthetic replicate, shown separately for primary reads (left) and high-quality reads (right). (B) Heatmaps of the mean percentage of reads mapped exclusively in the original dataset (“original-only”) or exclusively in the synthetic replicate (“replicate-only”), shown for primary reads (left) and high-quality reads (right).

## Supplementary Tables

**Table S1:**
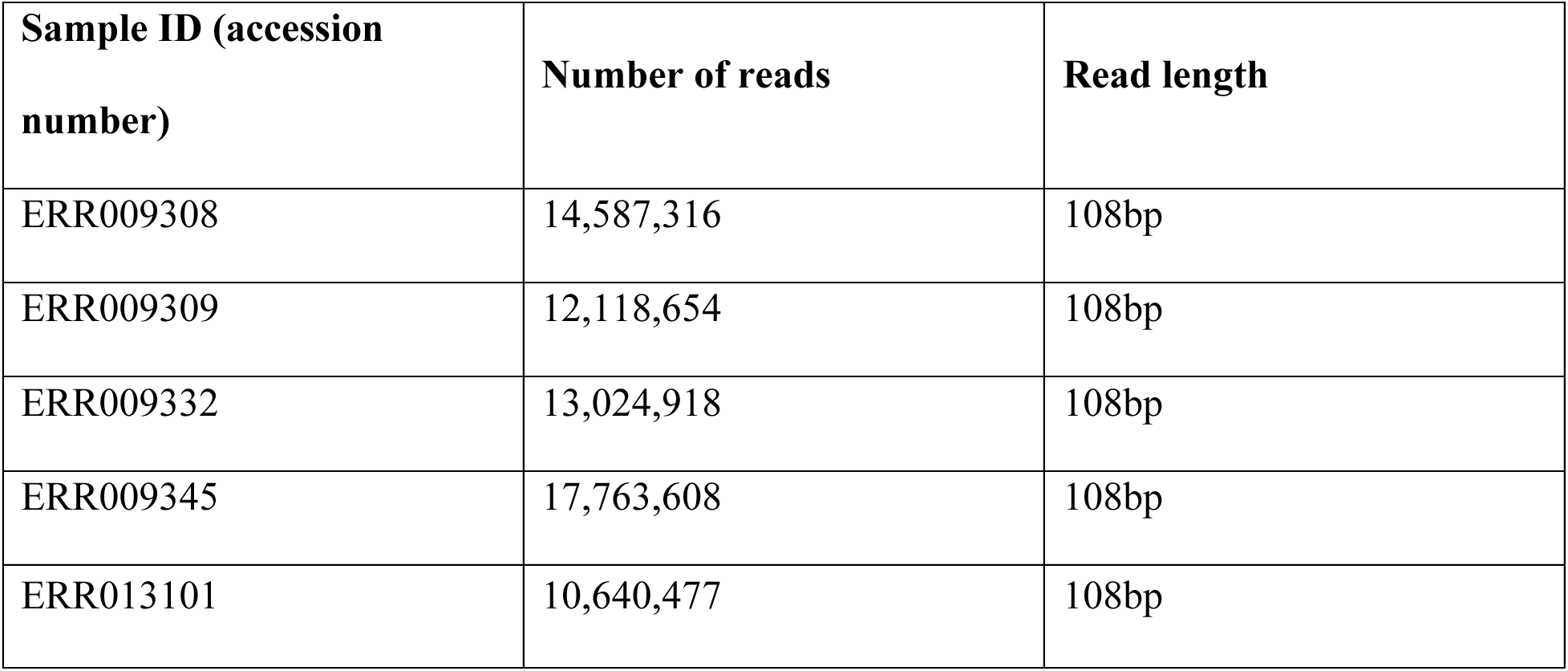
Selected WGS samples from the 1000 Genomes Project.

**Table S2:**
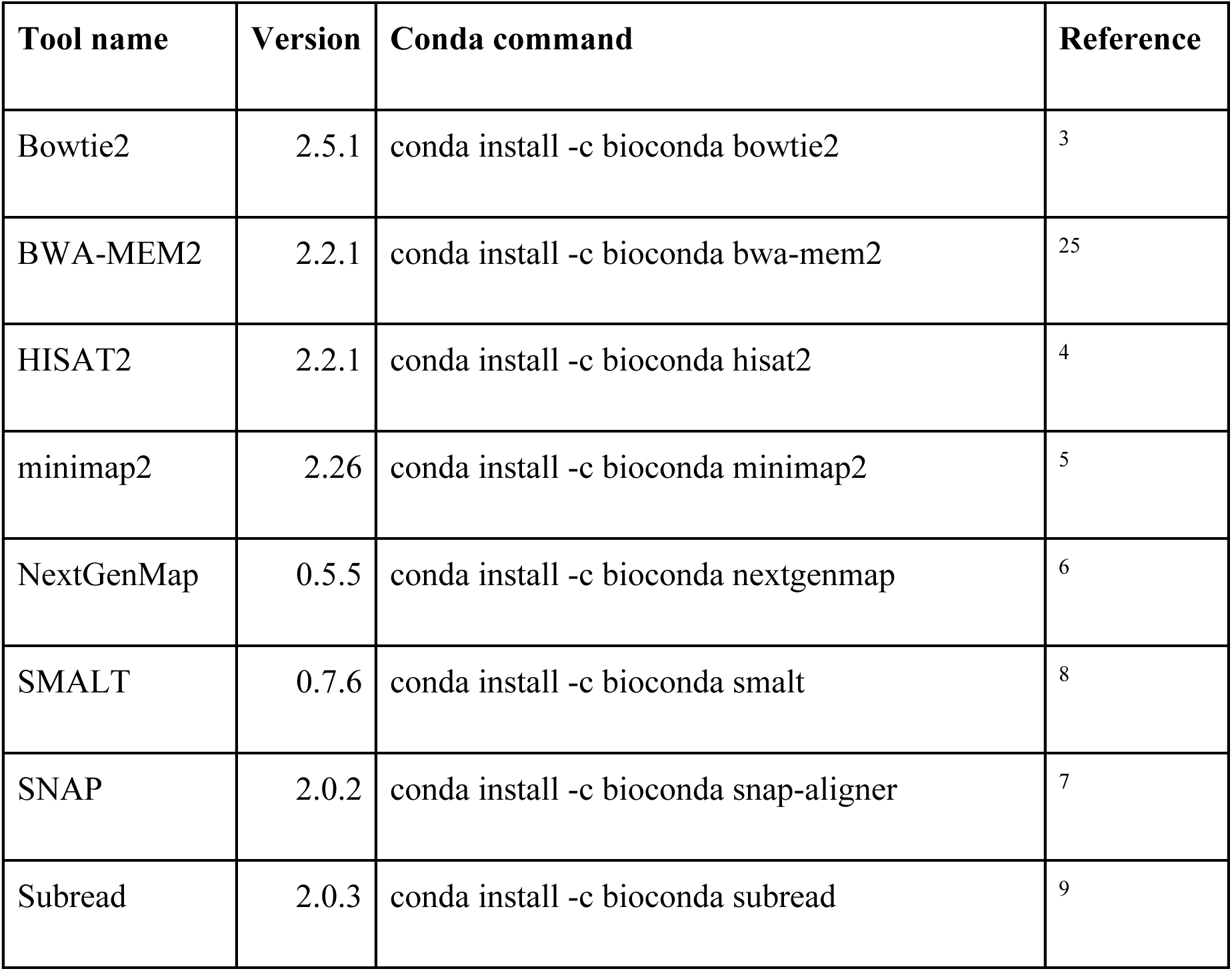
Overview of evaluated WGS read alignment tools.

**Table S3:**
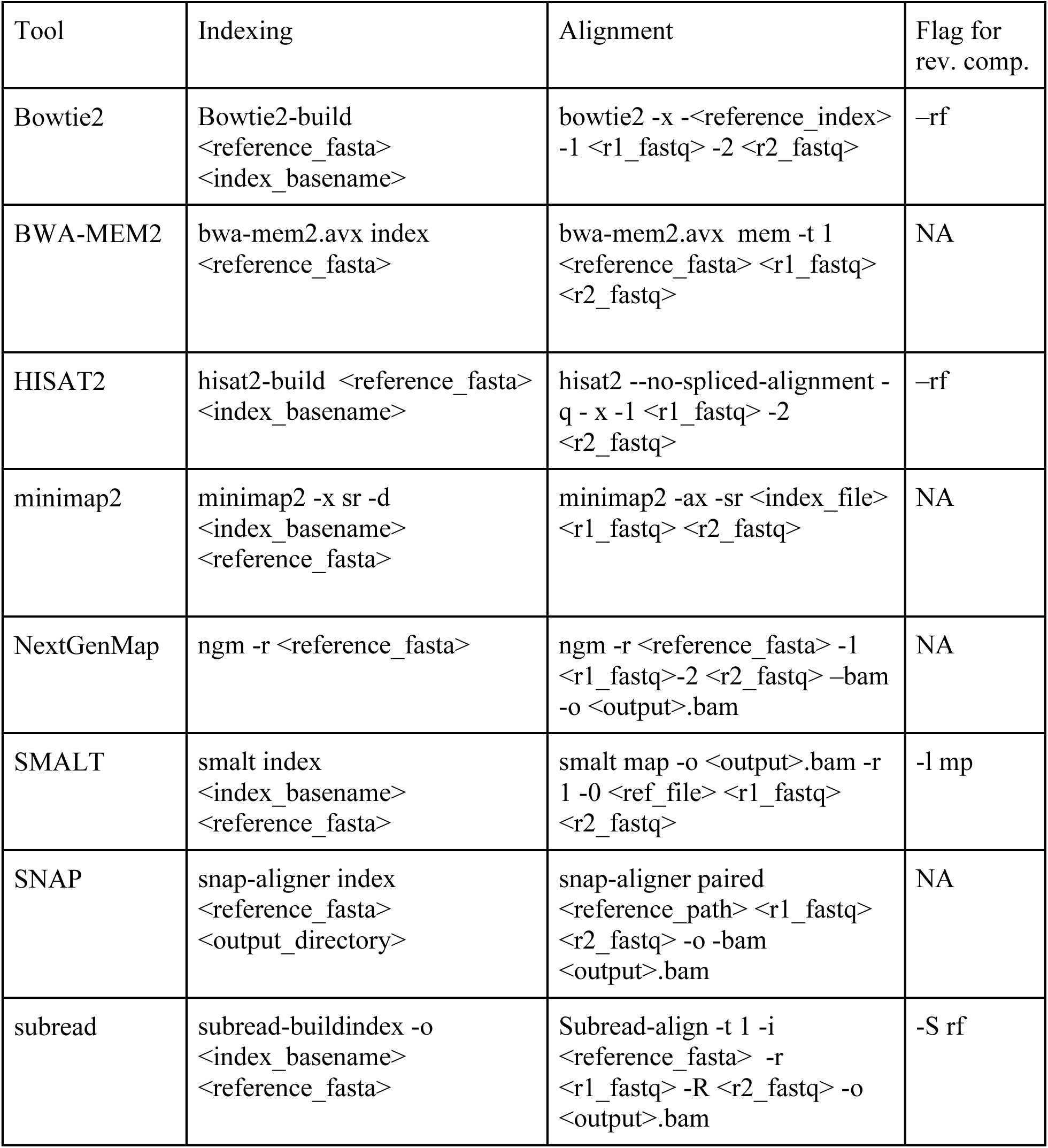
Command-line settings used for alignment reproducibility analysis.

**Table S4:**
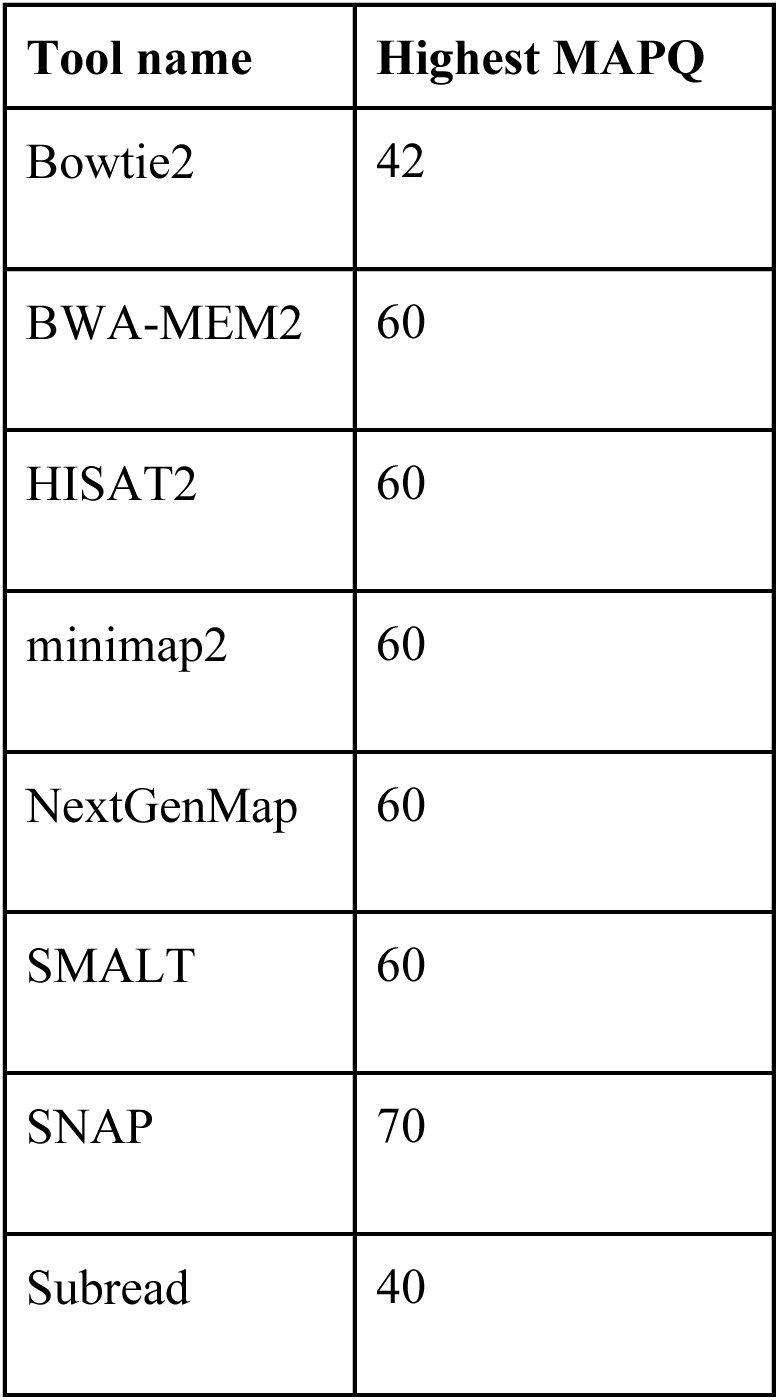
Maximum mapping quality (MAPQ) scores used to define high-quality reads for each tool.

| Tool name | Highest MAPQ |
| --- | --- |
| Bowtie2 | 42 |
| BWA-MEM2 | 60 |
| HISAT2 | 60 |
| minimap2 | 60 |
| NextGenMap | 60 |
| SMALT | 60 |
| SNAP | 70 |
| Subread | 40 |

**Table S5:** Summary of reproducibility classifications for read alignment tools across synthetic replicates. Primary reads and high-quality primary reads were evaluated separately. Columns represent synthetic replicate types (shuffling, reverse complementing, combined perturbation). Rows are divided into commonality and consistency metrics. “Full” under commonality indicates that 100% of mapped reads are shared between the original and replicate alignments, and 100% under consistency indicates identical mapping positions and edit distances among those shared reads. “Partial” denotes any value below 100% for either commonality or consistency. Lower percentages therefore reflect reduced commonality or alignment discrepancies. The table labels are based on numbers from figure 3 and 4.

| Primary reads | Shuffling |  | Reverse complementing |  | Combined perturbation |  |
| --- | --- | --- | --- | --- | --- | --- |
|  | commonality | consistency | common | consistency | commonality | consistency |
| Bowtie2 | Full | Full | Partial | Partial | Partial | Partial |
| BWA-MEM2 | Partial | Partial | Partial | Partial | Partial | Partial |
| HISAT2 | Full | Full | Partial | Partial | Partial | Partial |
| minimap2 | Full | Full | Partial | Partial | Partial | Partial |
| NextGenMap | Partial | Partial | Partial | Partial | Partial | Partial |
| SMALT | Partial | Partial | Partial | Partial | Partial | Partial |
| Snap | Full | Partial | Partial | Partial | Partial | Partial |
| Subread | Partial | Partial | Full | Full | Partial | Partial |

| High quality primary reads | Shuffling |  | Reverse complementing |  | Combined perturbation |  |
| --- | --- | --- | --- | --- | --- | --- |
|  | commonality | consistency | commonality | consistency | commonality | consistency |
| Bowtie2 | Full | Full | Partial | Partial | Partial | Partial |
| BWA-MEM2 | Partial | Partial | Partial | Partial | Partial | Partial |
| HISAT2 | Full | Full | Partial | Partial | Partial | Partial |
| minimap2 | Full | Full | Partial | Partial | Partial | Partial |
| NextGenMap | Partial | Full | Partial | Full | Partial | Full |
| SMALT | Full | Full | Partial | Partial | Partial | Partial |
| Snap | Partial | Full | Partial | Partial | Partial | Partial |
| Subread | Partial | Partial | Full | Full | Partial | Partial |

