## Supplemental Table 1 for "Assessing genomic reproducibility of read alignment tools"

| **Sample ID (accession number)** | **Number of reads** | **Read length** |
| --- | --- | --- |
| ERR009308 | 14,587,316 | 108bp |
| ERR009309 | 12,118,654 | 108bp |
| ERR009332 | 13,024,918 | 108bp |
| ERR009345 | 17,763,608 | 108bp |
| ERR013101 | 10,640,477 | 108bp |
