## Supplemental Table 2 for "Assessing genomic reproducibility of read alignment tools"

| **Tool name** | **Version** | **Conda command** | **Reference** |
| --- | --- | --- | --- |
| Bowtie2 | 2.5.1 | conda install -c bioconda bowtie2 | ^3^ |
| BWA-MEM2 | 2.2.1 | conda install -c bioconda bwa-mem2 | ^25^ |
| HISAT2 | 2.2.1 | conda install -c bioconda hisat2 | ^4^ |
| minimap2 | 2.26 | conda install -c bioconda minimap2 | ^5^ |
| NextGenMap | 0.5.5 | conda install -c bioconda nextgenmap | ^6^ |
| SMALT | 0.7.6 | conda install -c bioconda smalt | ^8^ |
| SNAP | 2.0.2 | conda install -c bioconda snap-aligner | ^7^ |
| Subread | 2.0.3 | conda install -c bioconda subread | ^9^ |
