## Supplemental Table 3 for "Assessing genomic reproducibility of read alignment tools"

| Tool | Indexing | Alignment | Flag for rev. comp. |
| --- | --- | --- | --- |
| Bowtie2 | Bowtie2-build <reference_fasta> <index_basename> | bowtie2 -x -<reference_index> -1 <r1_fastq> -2 <r2_fastq> | –rf |
| BWA-MEM2 | bwa-mem2.avx index <reference_fasta> | bwa-mem2.avx  mem -t 1 <reference_fasta> <r1_fastq> <r2_fastq> | NA |
| HISAT2 | hisat2-build  <reference_fasta> <index_basename> | hisat2 --no-spliced-alignment -q - x -1 <r1_fastq> -2 <r2_fastq> | –rf |
| minimap2 | minimap2 -x sr -d <index_basename> <reference_fasta> | minimap2 -ax -sr <index_file> <r1_fastq> <r2_fastq> | NA |
| NextGenMap | ngm -r <reference_fasta> | ngm -r <reference_fasta> -1 <r1_fastq>-2 <r2_fastq> –bam -o <output>.bam | NA |
| SMALT | smalt index <index_basename> <reference_fasta> | smalt map -o <output>.bam -r 1 -0 <ref_file> <r1_fastq> <r2_fastq> | -l mp |
| SNAP | snap-aligner index <reference_fasta> <output_directory> | snap-aligner paired <reference_path> <r1_fastq> <r2_fastq> -o -bam <output>.bam | NA |
| subread | subread-buildindex -o <index_basename> <reference_fasta> | Subread-align -t 1 -i <reference_fasta>  -r <r1_fastq> -R <r2_fastq> -o <output>.bam | -S rf |
