## Supplementary figures and images for "Assessing genomic reproducibility of read alignment tools"

### Supplemental Table 5

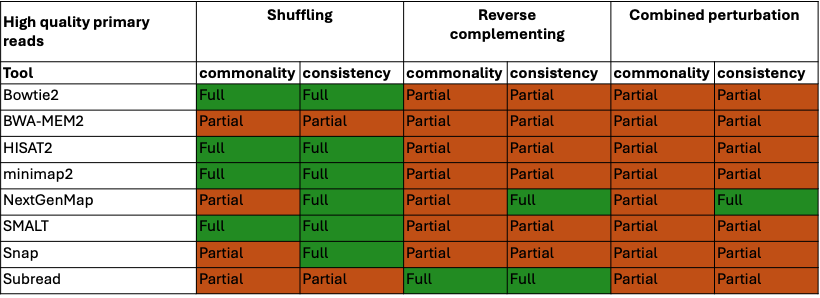

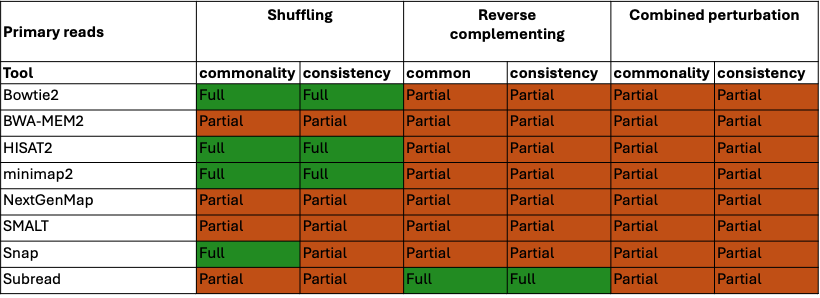
